# CD61 identifies a superior population of aged murine hematopoietic stem cells and is required to preserve quiescence and self-renewal

**DOI:** 10.1101/2023.08.14.553203

**Authors:** Natalia Skinder, Irene Sanz Fernández, Albertien Dethmers-Ausema, Ellen Weersing, Gerald de Haan

## Abstract

Aging leads to a decline in function of hematopoietic stem cells (HSCs) and increases susceptibility to hematological disease. We found CD61 to be highly expressed in aged HSCs. Here we investigate the role of CD61 in identifying distinct subpopulations of aged HSCs and assess how expression of CD61 affects stem cell function. We show that HSCs with high expression of CD61 are functionality superior and retain self-renewal capacity in serial transplantations. A population of aged HSCs with highest CD61 expression is functionally comparable to young HSCs. These CD61^High^ HSCs display a notably higher quiescence compared to their CD61^Low^ counterparts. We also show that CD61^High^ and CD61^Low^ HSCs are transcriptomically distinct populations within aged HSCs. Collectively, our research identifies CD61 as a key player in maintaining stem cell quiescence during aging, ensuring the preservation of their functional integrity and potential. Moreover, CD61 emerges as a marker to prospectively isolate a superior, highly dormant population of young and aged HSCs, making it a valuable tool both in fundamental and clinical research.

**Highlights:**

- CD61 expression marks a functionally superior population of long-term hematopoietic stem cells and plays a pivotal role in the maintenance of aged LT-HSCs by regulating their dormancy
- Aged LT-HSCs with high expression of CD61 are functionally comparable to their young counterparts

## Introduction

Hematopoietic stem cells (HSCs) reside at the top of the hematopoietic hierarchy and are defined by their unique regenerative potential and ability to give rise to all the blood lineages throughout the lifetime of an organism. Nevertheless, normal aging has been shown to lead to a gradual functional decline of the hematopoietic stem cells^1–3^. Upon aging, murine HSCs lose their overall self-renewal and regenerative potential, skew towards the myeloid lineage and expand in numbers^4–7^.

Long-term transplantation assays show that aged HSCs, when transplanted into a young bone marrow (BM) microenvironment, display a reduced functional activity, hence the age-related decrease in HSC activity seems to be mainly intrinsically-driven^8^. Numerous cell-intrinsic defects have been suggested to contribute to the decreased overall potential of aged HSCs, such as the loss of polarity^9,10^, impaired autophagy^11,12^, DNA damage accumulation^13,14^ and epigenetic modifications^15,16^. However, it has also been shown that an altered BM microenvironment affects the functional activity of aged of HSCs. For example, aged HSCs transplanted into young recipients exhibited reduced myeloid-skewing compared to those transplanted into old recipients^17^. In addition, BM niche remodeling and localization of HSCs in their niche affect the aging process^18,19^. HSC aging is thus a very heterogenous process influenced and regulated by numerous intrinsic and extrinsic mechanisms, where not every individual HSC is equally susceptible to or affected by the aging process ^20–22^. Interestingly, some individual aged HSCs have been shown to function as if they have not aged at all^4,6^.

Recently we published a meta-analysis of 16 independent studies in which the transcriptomes of young and aged murine HSCs had been assessed. From these studies we were able to derive an HSC Aging Signature – a robust list of genes that are differentially expressed upon murine HSC aging^23^. This signature, consisting of ∼200 genes, included an abundance of novel genes previously not associated with aging, nor with HSCs. Unexpectedly, almost 50% of the genes in the Aging Signature encode for membrane-associated genes, which suggests that communication between HSCs and the bone marrow environment is altered during aging.

Itgb3, a membrane-associated gene that encodes for CD61 – a widely expressed β-integrin from the integrin family^24,25^, was upregulated in 9 out 12 studies used to establish the Aging Signature. CD61 acts in heterodimers with the α-integrins CD41 and CD51, and is known to regulate cell adhesion, cell signaling and differentiation^26–28^. Notably, it is the only member of the integrin family to be present in the Aging Signature. The role of CD61 in regulating HSCs has been studied previously and CD61 has been reported to affect several HSC-niche communication pathways, including thrombopoietin-mediated HSC regulation^29^, cell adhesion and inflammation response via interferon γ signaling^30^. High expression of CD61 has also been shown to be correlated with a more quiescent subpopulation of HSCs in young mice^31^. Notwithstanding these effects, it has also been shown that downregulation or deletion of CD61 in HSCs had little to no effect on the overall functionality of HSCs^29,32^. CD61 has never been studied in the context of HSC aging.

As we found CD61 to be one of the most robustly upregulated genes in aged HSCs, we aimed to assess whether altered expression of CD61 on aged HSCs affects stem cell functioning. Our study shows that CD61 expression identifies a functionally superior population of LT-HSCs in both young but particularly aged bone marrow and may be used to prospectively isolate these cells.

## Materials and Methods

### Mice

All experiments were approved by the Central Commission for animal Testing and Animal Ethical Committee. Young (2-4 months), middle-aged (10-12 months) and aged (>22 months) C57BL/6J were obtained from either CDP or Janvier labs, France. All mice were housed in a temperature and day cycle-controlled conditions.

### Flow Cytometry

Bone marrow was isolated from long bones, erythrocytes were lysed. For cell isolation, the lysed bone marrow cells were stained with antibodies used to detect stem cells and PI for Live/Dead staining. Cells were isolated on MoFlo Astrios or MoFlo XDP cell sorters.

### In vitro experiments

#### Single cell colony assay

Single LT-HSCs were sorted and cultured for 14 days in StemSpan supplemented with 100U/ml penicillin, 100μg/ml streptomycin, 10% Australian FCS, 300ng/ml SCF, 20ng/ml IL-11 and 1ng/ml Flt3 ligand. The size of the colonies was analyzed after 7 and 14 days.

#### Single cell division assay

Single LT-HSCs were sorted and cultured for 2 days in StemSpan supplemented with cytokines (described above). The number of cells was analyzed after 48h.

#### *γ*H2AX immunofluorescence staining

1.000-2.000 LT-HSCs were sorted an adhesion slide. Cells were then fixed, permeabilized and blocked. Cells were stained with primary α-γH2AX and secondary conjugated antibody and the coverslip was mounted. The slide was imaged on a Leica Sp8 confocal microscope. The data were analyzed using Fiji Image J.

#### LT-HSC cell cycle analysis

LT-HSCs were sorted, fixed and stained according to the manufacturer’s protocol. Cells were stained with α-Ki67 antibody and DAPI. Samples were analyzed on BD FACS Canto II or BD Symphony.

#### Inhibitor treatment of LT-HSCs

The inhibitor treatment was done in single-cell proliferation assay, division assay, cell cycle analysis according to the protocol described above with medium supplemented with 50ng/mL cycloRGDγK, 25ng/mL Tirofiban or DMSO.

For single-cell expansion assays, HemEx-Type9A medium was supplemented with 100ng/ml TPO, 10ng/ml SCF, 50ng/mL cycloRGDγK, 25ng/mL Tirofiban or DMSO.

#### Quantitative PCR analysis

RNA was isolated by using RNeasy Micro Kit. The RNA was transcribed using SuperScript VILO cDNA Synthesis Kit. The amplicons for CD61 and housekeeping gene Gapdh were amplified and quantified via qPCR LightCycler 480 Instrument.

### Transplantation

#### CD61^High^ and CD61^Low^ LT-HSC transplantation

CD61^High^ and CD61^Low^ LT-HSCs were transplanted together with 2 x 10^6^ competitor W^41^ mice (C57BL/6J-KitW-41J/J) BM cells into gender-matched, lethally irradiated (9 Gy) recipients. 16 weeks after transplantation, donor-derived LT-HSCs were transplanted alongside 22 x 10^6^ competitor W^41^ mice (C57BL/6J-KitW-41J/J) BM cells into gender-matched lethally irradiated (9 Gy) secondary recipients.

#### CD61^KD^ LT-HSC transplantation

##### Transduction

LT-HSCs were isolated and plated 24h prior to transduction in HemEx-Type9A medium supplemented with 100ng/ml TPO and 10ng/ml SCF. The viral supernatant was added to the LT-HSCs next day.

##### Transplantation

After 5 days mCherry^+^ LT-HSCs were transplanted alongside 2 million CD45.1^+^ W41 BM cells into gender-matched lethally irradiated (9 Gy) recipients.

#### Peripheral blood count

Peripheral blood was collected from the retro-orbital venous plexus in heparinized capillary tubes. 25 μl PB was used for cell counting using Medonic CA-620.

### RNA-sequencing

Total RNA was isolated from 5000 LT-HSCs using RNeasy Plus Micro Kit (QIAGEN, 74034) according to the manufacturer’s instructions. Library preparation was performed with SMART Ultra-Low Input kit v4 (Takara) and Nextera™ XT Library Prep kit (Illumina). Samples were sequenced using a NextSeq 5000 (Illumina) in the same flow cell and were pulled equimolarly.

Data analysis FASTQ files were quality-control checked using FastQC (0.11.9) and Picard (2.23.0). Following, reads were mapped to reference mouse genome (GENCODE, GRCm38, M21) using STAR (2.7.0d) with standard arguments and --outFilterMatchNminOverLread 0.4--outFilterScoreMinOverLread 0.4. Unstranded read counts were used to perform differential expression analysis using DESeq2.

### Statistical analysis

All experiments were performed as least 2 times. The number of mice or technical replicates is indicated in the figure legends. Data are shown as mean ± SD. Unpaired, two-tailed Student t-test and two-way ANOVA with Sidak’s multiple comparison test were performed in GraphPad Prism 9.0 and 10.0. Significant p-value was indicated as *p<0.05, **p<0.01, ***p<0.001 and ****p<0.0001.

### Data Sharing Statement

RNA-seq data was deposed under the accession number GSE240650.

## Results

### During aging expression of CD61 is specifically increased in the most primitive hematopoietic stem cells

Recently, we published a transcriptome meta-analysis which revealed that Itgb3 – the gene encoding for CD61 - is the only consistently reported integrin family member that is differentially expressed upon murine hematopoietic stem cell aging (Fig 1A). Using qRT-PCR we analyzed various subpopulations within the progenitor cell compartment (Lin^-^, Sca1^+^, c-Kit^+^) and found that Itgb3 is exclusively upregulated in the most primitive LT-HSCs (Lin^-^, Sca1^+^, c-Kit^+^, CD150^+^, CD48^-^) (Fig. 1B). With FACS we confirmed that increased Itgb3 mRNA expression in aged LT-HSCs correlates with increased protein levels (Fig. 1C-D).

**Figure 1.**
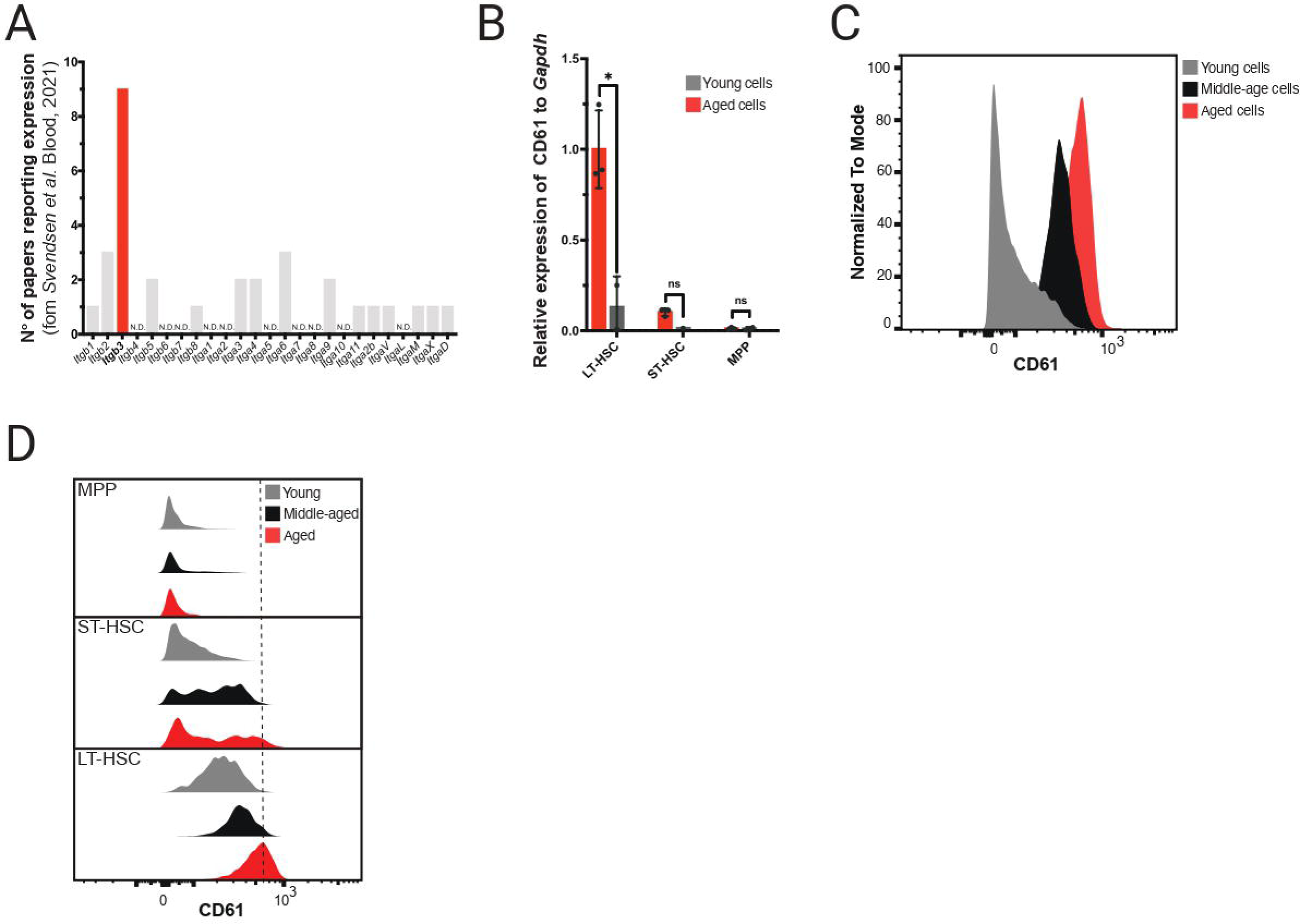
-CD61 expression in LT-HSCs. **A** – Number of papers in Aging Signature reporting differential expression of genes of the integrin family **B** – CD61 mRNA expression measured by RT_PCR in LT-HSCs, ST-HSCs and MPP isolated from young and aged mice **C** - Protein level of CD61 on LT-HSCs from young, middle-aged and aged mice measured by flow cytometry **D** - Protein level of CD61 on LT-HSCs, ST-HSCs and MPPs from young, middle-aged and aged mice measured by flow cytometry

Furthermore, to assess how expression of CD61 changes during the aging trajectory, we analyzed its expression at different ages. This revealed that CD61 expression increases gradually throughout the lifetime of a mouse (Fig.1C-D).

### Aged CD61^High^ and CD61^low^ express distinct transcriptomes

To better understand the role of CD61 upregulation in LT-HSCs upon aging, we performed RNA sequencing on highly purified aged LT-HSCs separated by CD61 expression. To this end we isolated the 10% highest (CD61^High^) and lowest (CD61^Low^) CD61 expressing LT-HSCs (Fig. 2A and B).

**Figure 2.**
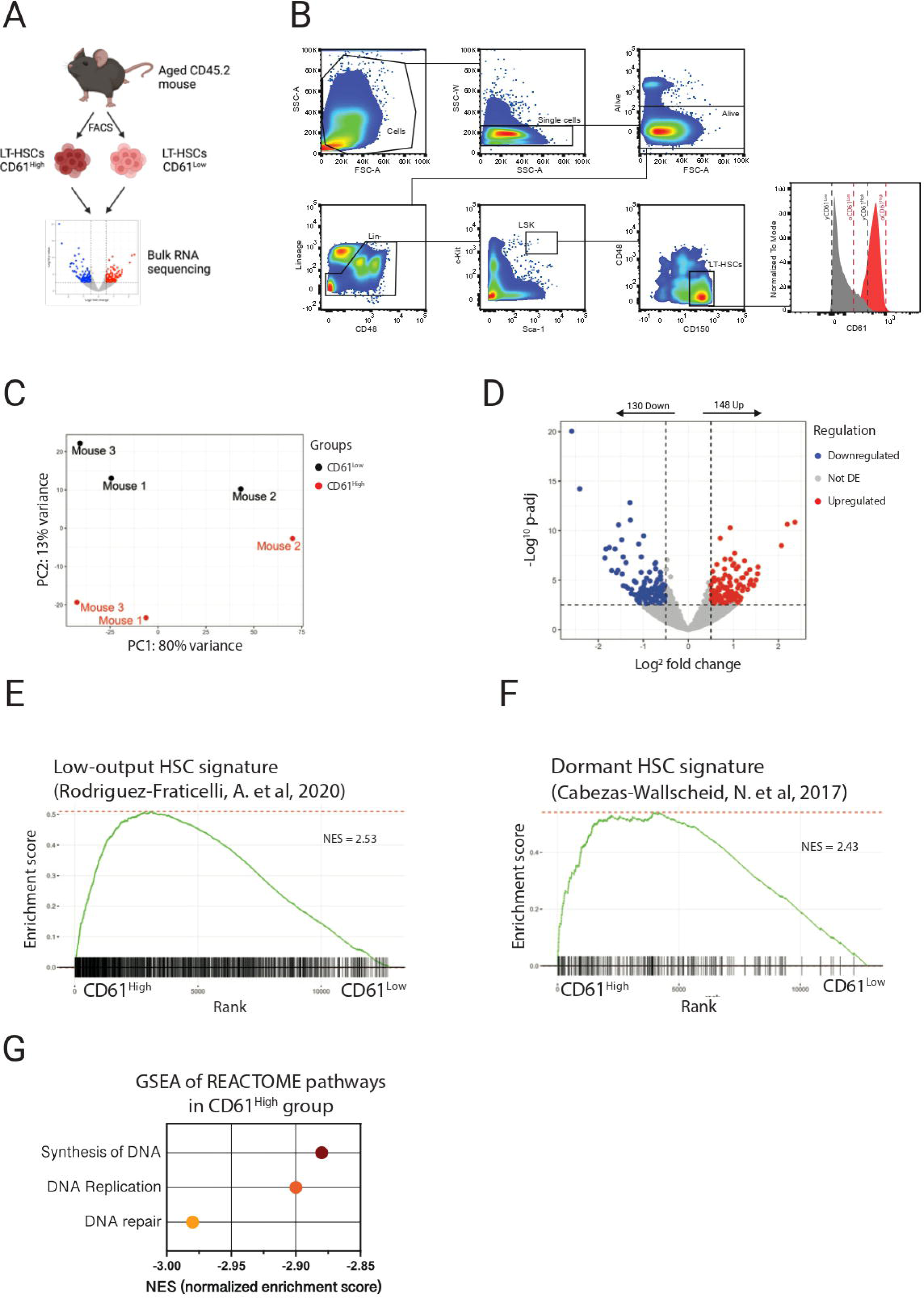
– Transcriptome analysis of aged CD61^high^ and CD61^low^ LT-HSCs. **A** – Schematic representation of the RNA sequencing experiment **B** – Sorting strategy for **CD61**^high^ and **CD61**^low^ LT-HSCs **C** – MDS plot showing the Principal Coordinates Analysis (PCA) of CD61^High^ and CD61^Low^ LT-HSCs **D** – DEGs in aged CD61^High^ and CD61^Low^ LT-HSCs. Volcano plot showing distribution of the adjusted p-value (-log_10_ p-value) and the fold changes (logFC). Up-regulated and down-regulated genes are indicated in red and blue, respectively (p-adj < 0.01) **E** – Signature enrichment plot from GSEA using Low-Output HSC gene set in LT-HSCs from CD61^High^ and CD61^Low^ cells (GSE134242) **F** - Signature enrichment plot from GSEA using Dormant HSC gene set in LT-HSCs from CD61^High^ and CD61^Low^ cells (GSE87814) **G** – GSEA for the most down-regulated pathways in CD61^High^ LT-HSCs

Even though Lin^-^Sca1^+^c-Kit^+^CD150^+^CD48^-^ are highly purified, we found that within this population CD61^High^ and CD61^Low^ expressing cells differed significantly in their transcriptomic landscape (Fig. 2C). We identified 278 genes that were significantly differentially expressed between CD61^High^ and CD61^Low^ LT-HSCs, with 148 upregulated and 130 downregulated genes in CD61^High^ LT-HSCs (Fig. 2D). To compare CD61^High^ and CD61^Low^ LT-HSC transcriptome signatures, we performed gene set enrichment analysis (GSEA) for previously published LT-HSC gene signatures (Rodriguez-Fraticelli, 2020; Cabezas-Wallscheid, 2017). Interestingly, CD61^High^ LT-HSCs showed enrichment for a dormant HSC signature, as well as low-output HSC signature (Fig 2E-F). Both low-output and dormant HSCs are stem cell populations associated with superior reconstitution and self-renewal potential^33,34^. Additionally, GSEA analysis showed enrichment of cell cycle-related pathways in CD61^Low^ LT-HSCs, suggesting that these cells are actively cycling, compared to CD61^High^ LT-HSCs (Fig. 2G). Together, these data show that aged CD61^High^ LT-HSCs are molecularly distinct from CD61^Low^ LT-HSCs and exhibit a signature that is associated with superior stem cell functionality.

### High expression of CD61 in aged LT-HSCs is associated with quiescence

The RNA sequencing data revealed that aged LT-HSCs with differential CD61 expression are molecularly distinct (Fig. 2C-D). We next tested whether these molecular differences would translate into functional consequences.

We first analyzed the proliferation potential of CD61^Low^ and CD61^High^ LT-HSCs, isolated from young or aged donor mice, by sorting single LT-HSCs in a well and culturing these for 14 days. We analyzed the size of the colony that they were able to produce. After 14 days young CD61^High^ and CD61^Low^ LT-HSCs proliferated equally (Fig. 3B). However, when isolated from aged mice, CD61^Low^ LT-HSCs proliferated significantly worse, with a large fraction of cells not proliferating at all (Fig. 3C). Aged CD61^High^ LT-HSCs proliferated similarly to cells isolated from young mice, confirming their “young-like” phenotype.

**Figure 3.**
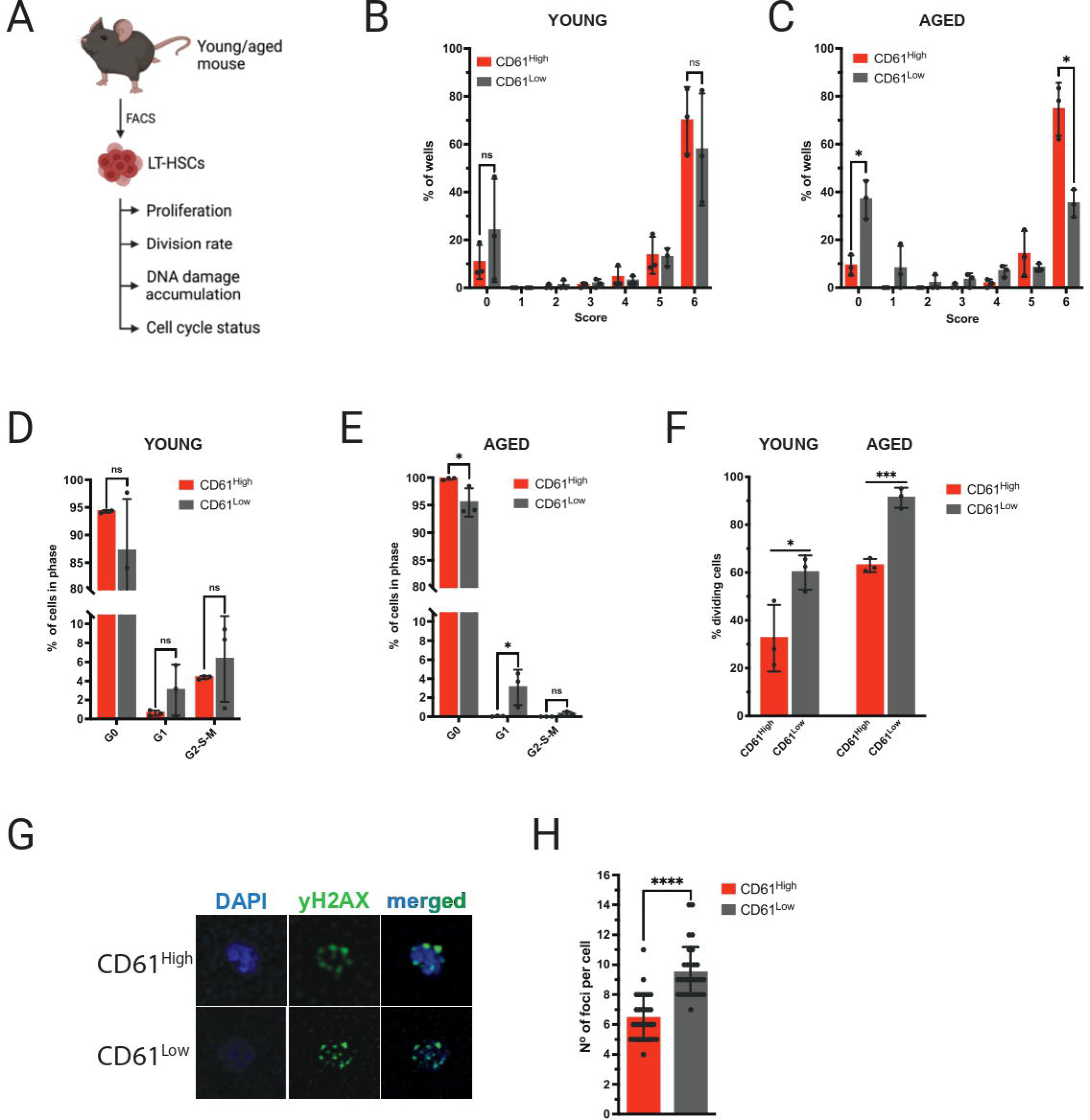
– In vitro experiments. **A** – Schematic representation of in vitro experiments performed with young and aged LT-HSCs **B** - Single cell proliferation assay of young CD61^High^ and CD61^Low^ LT-HSCs; Size 0: no cells; Size 1: 1-30 cells; Size 2: 31-100 cells; Size 3: 101-1000 cells; Size 4: 5000 cells; Size 5: 15000 cells Size 6: 30000 cells; Size 7: 30000< cells **C** - Single cell proliferation assay of aged CD61^High^ and CD61^Low^ LT-HSCs; **D** - Cell cycle analysis using Ki67/DAPI staining of young CD61^High^ and CD61^Low^ LT-HSCs E - Cell cycle analysis using Ki67/DAPI staining of aged CD61^High^ and CD61^Low^ LT-HSCs F - Single cell division assay of young and aged CD61^High^ and CD61^Low^ LT-HSCs **G** – DAPI/γH2AX immunofluorescent staining of aged CD61^High^ LT-HSCs, CD61^Low^ LT-HSCs and LT-HSCs **H** – Number of γH2AX foci per cell in CD61^High^ and CD61^Low^ aged LT-HSCs

As the transcriptome data suggested that CD61^High^ and CD61^Low^ LT-HSCs may differ in their cell cycle activity, we performed cell cycle analysis using Ki67 and DNA staining. This experiment showed that young CD61^High^ and CD61^Low^ LT-HSCs had a comparable distribution of cells in distinct cell cycle phases (Fig. 3D). In contrast, in aged LT-HSCs essentially all CD61^High^ LT-HSCs were in G_0_, whereas CD61^Low^ cells were more abundant in G1 and G2-S-M phases (Fig. 3E).

Next, we analyzed the division rate of young and aged CD61^High^ and CD61^Low^ LT-HSCs, by sorting single-cell into a well and after 48h counting the fraction of wells in which a cell division had occurred, i.e., in which more than one cell was present. For both young and aged we found a lower division rate of CD61^High^ LT-HSCs (Fig. 3F). These functional data corroborate the molecular data and show that CD61^High^ LT-HSCs are more quiescent.

### LT-HSCs with low expression of CD61 accumulate significantly more DNA damage

Hypothesizing that the higher proliferation rate of aged CD61^Low^ LT-HSCs may result in increased DNA damage accumulation and subsequent cell death, we analyzed levels of DNA damage in aged stem cells. We stained freshly isolated aged CD61^High^ and CD61^Low^ LT-HSCs with an anti-γH2AX antibody, which detects double-strand breaks (DSB) within cells. Using confocal microscopy, we imaged cells and counted the number of γH2AX foci. Figure 3G shows a representative comparison between aged CD61^High^ and CD61^Low^ cells. The analysis revealed that CD61^Low^ LT-HSCs exhibited a significantly higher number of foci per cell compared to CD61^High^ cells (Fig. 3H). This confirms that aged LT-HSCs with low CD61 expression accumulate more DNA damage, likely due to their higher division rate and proliferative potential. Consequently, increased DNA damage may trigger apoptosis, providing a possible explanation for their lower output in in vitro single-cell proliferation assays.

### CD61-CD51 dimerization drives CD61-mediated maintenance of aged LT-HSCs quiescence

Integrins dimerize into distinct hetero- and homodimers to facilitate biological processes. CD61 can heterodimerize with CD51 (Itgav) or CD41 (Itga2b)^35^. To further understand the mechanism of CD61-mediated LT-HSCs quiescence we explored which CD61 heterodimer is involved in this process.

We first determined whether either CD51 or CD41 is co-expressed with CD61 in aged LT-HSCs. As shown in Figure 4B, CD51, but not CD41, expression levels correlated with CD61 expression. This suggests that CD61-CD51 dimerization may occur in aged LT-HSCs.

**Figure 4.**
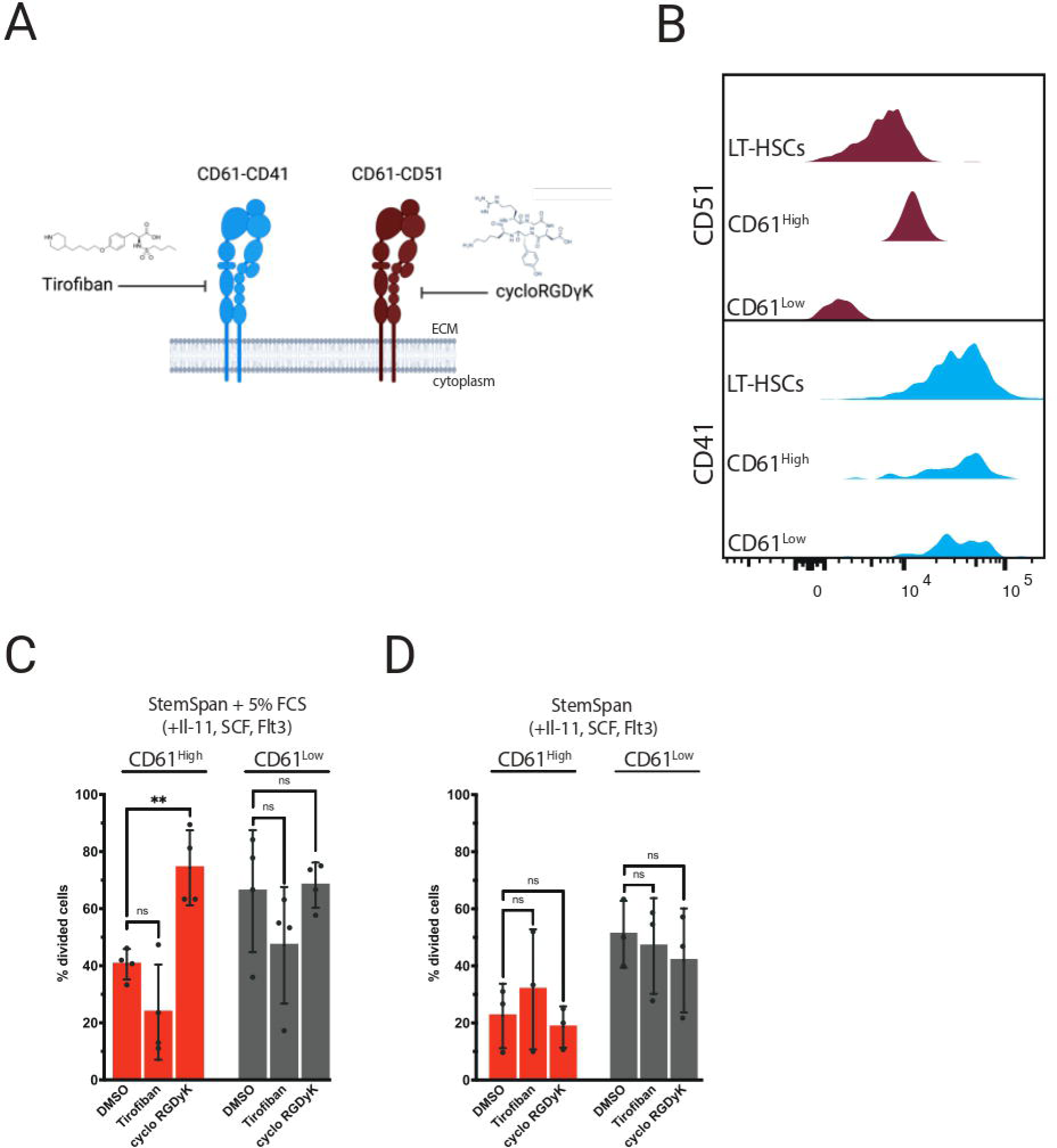
– Inhibitor studies. **A** – Schematic representation of inhibition of CD61 signaling with small molecules **B** – Histogram representing co-expression of CD51 or CD41 with CD61 in aged LT-HSCs **C** – Single cell division assay of aged CD61^High^ and CD61^Low^ LT-HSCs treated with Tirofiban or cycloRGDγK in StemSpan with 5% serum supplementation and cytokines **D** - Single cell division assay of aged CD61^High^ and CD61^Low^ LT-HSCs treated with Tirofiban or cycloRGDγK in serum-free StemSpan supplemented with cytokines

We then cultured aged LT-HSCs in the presence of dimer-specific small-molecule inhibitors – Tirofiban for CD61-CD41 and cycloRGDγK for CD61-CD51 (Fig 4B). Both inhibitors have been reported to selectively inhibit the activity of their target dimer^36,37^. We analyzed whether these inhibitors affected proliferation kinetics of aged CD61^High^ and CD61^Low^ LT-HSCs. As expected, Tirofiban had a minimal effect, whereas cycloRGDγK supplementation induced increased cycling in LT-HSCs in serum-containing cultures (Fig. 4C). Interestingly, this effect was restricted to (quiescent) CD61^High^ stem cells, increasing their division rate comparable to those of CD61^Low^ LT-HSCs. The release of quiescence was not detectable in unfractionated LT-HSCs, which suggests that CD61-CD51 heterodimerization occurs specifically in LT-HSCs with highest CD61 expression (Supplementary Fig. S4A). Cell cycle analysis confirmed that CD61^High^ LT-HSCs treated with cycloRGDγK had a higher percentage of cells in G2-S-M phases in comparison to DMSO control. In addition, we found a higher number of cells in cycloRGDγK-treated cultures (Supplementary Fig. S4C).

### CD61 expression marks a subpopulation of functionally superior aged LT-HSCs

The transcriptomic data as well as in vitro experiments indicated that CD61 expression marks distinct populations of LT-HSCs. We next asked whether these differences would also translate in functional differences in in vivo assays. To this end we employed competitive transplantation assays using freshly isolated CD61^High^ and CD61^Low^ LT-HSCs (CD45.2^+^). We transplanted 500 CD61^High^ and 500 CD61^Low^ LT-HSCs, along with 2×10^6^ whole bone marrow cells (WBM) obtained from W^41^ mice (C57BL/6J-KitW-41J/J), into lethally irradiated recipient mice (CD45.1^+^). Chimerism levels in the peripheral blood (PB) of recipient mice were analyzed every 4 weeks for at least 16 weeks using flow cytometry. After 16 weeks recipient mice were sacrificed and their bone marrow analyzed. Next, 500 LT-HSCs harvested from primary recipients were serially competitively transplanted into secondary recipients and chimerism levels were analyzed for another 16 weeks. After the second 16-week period bones were harvested and analyzed.

We first competitively transplanted CD61^High^ and CD61^Low^ LT-HSCs isolated from young mice. In young LT-HSCs, differential expression of CD61 did not affect the reconstituting potential in primary transplantations (Fig 5B, 5D). CD61^High^ and CD61^Low^ LT-HSCs isolated from young mice possessed comparable lineage contribution to the myeloid, T-cell and B-cell peripheral blood cell compartments (Supplementary Fig. S5J). However, when we performed these experiments using aged mice as donors, we observed significant differences in chimerism levels between mice transplanted with CD61^High^ or CD61^Low^ LT-HSCs as early as 4 weeks post-transplantation (Fig 5C-D).

**Figure 5.**
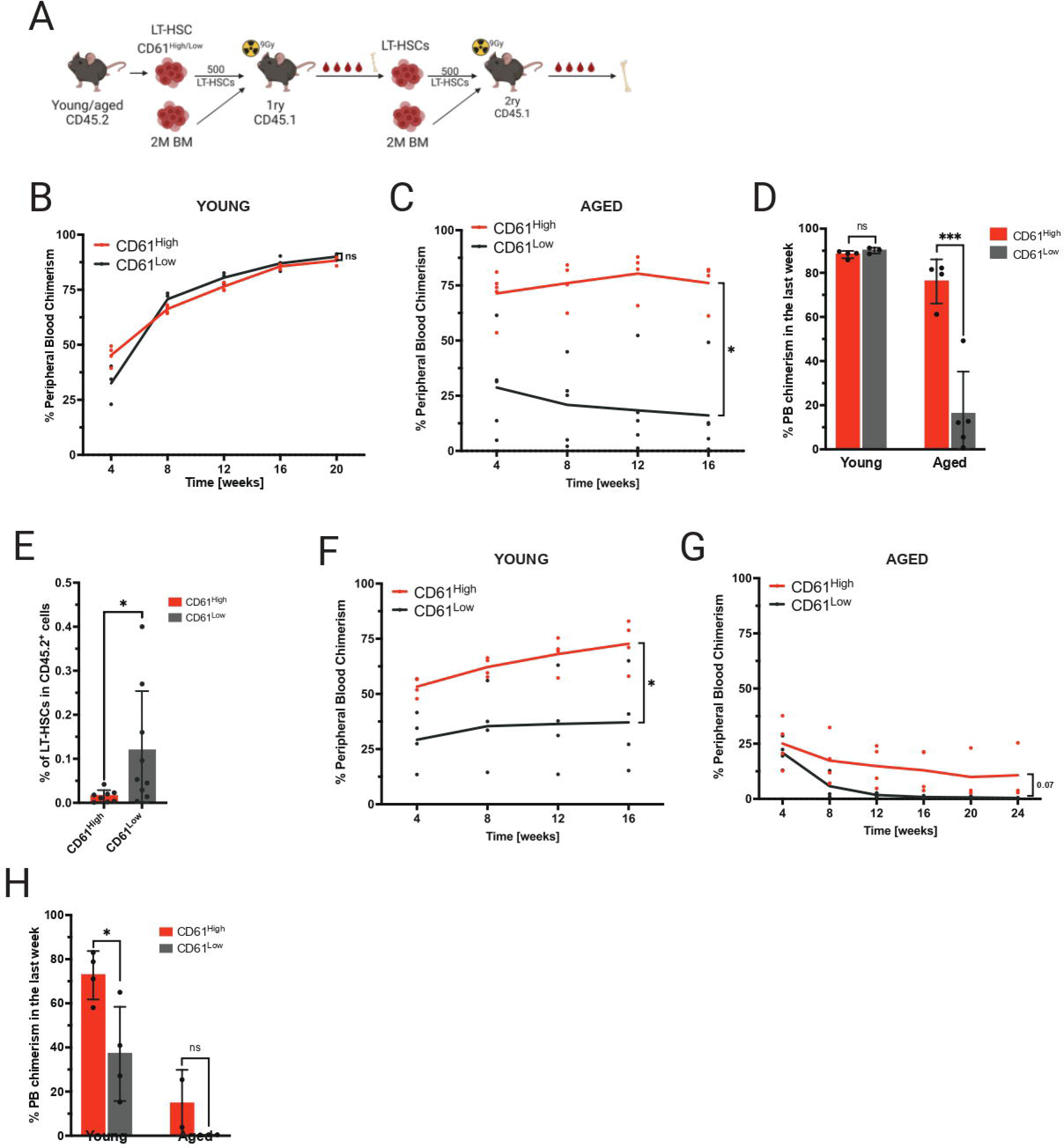
– Transplantation studies. **A** – Schematic representation of competitive primary and secondary transplantation assays **B** –Competitive transplantation assay of 500 CD61^High^ and CD61^Low^ LT-HSCs from young mice into lethally irradiated recipients (mice per group – n=4). Data from representative experiment are shown **C** – Competitive transplantation assay of 500 CD61^High^ and CD61^Low^ LT-HSCs from aged mice into lethally irradiated recipients (mice per group – n=4). Data from representative experiment are shown. **D** - Peripheral blood chimerism levels long-term post-transplantation in primary recipients of young and aged CD61^High^ and CD61^Low^ LT-HSCs **E** – Donor-derived LT-HSC frequency in BM of primary recipients transplanted with aged donor CD61^High^ and CD61^Low^ LT-HSCs **F** – Secondary transplantation assay of 500 LT-HSCs derived from primary recipients transplanted with young CD61^High^ and CD61^Low^ LT-HSCs (mice per group – n=4). Data from representative experiment are shown. **G** - Secondary transplantation assay of 500 LT-HSCs derived from primary recipients transplanted with aged CD61^High^ and CD61^Low^ LT-HSCs (mice per group – n=4). Data from representative experiment are shown. **H** - Peripheral blood chimerism levels long-term post-transplantation in secondary recipients of young and aged CD61^High^ and CD61^Low^ LT-HSCs

Aged CD61^Low^ LT-HSCs produced significantly less peripheral blood cells compared to CD61^High^ LT-HSCs. Chimerism levels in mice transplanted with aged CD61^Low^ LT-HSCs remained below 25% whereas contribution derived from aged CD61^High^ cells gradually increased, and were comparable to young HSCs (5B, C). The lineage contribution to different PB cell populations was similar between CD61^High^ and CD61^Low^ LT-HSCs (Supplementary Fig. S5M).

After 16 weeks the BM of primary recipient mice was analyzed and although no significant differences were found in the bone marrow of recipient mice transplanted with young CD61^High^ or CD61^Low^ LT-HSCs, we discovered an average 8-fold increase in the frequency of the donor-derived aged CD61^Low^ LT-HSCs (Fig. 5E). As HSCs highly rely on their self-renewal/differentiation ratio and its balance to maintain their activity throughout the lifetime of the organism, we hypothesized that the reason for this increase is a higher proliferation rate of cells with a decreased CD61 expression.

We performed secondary transplantations to determine self-renewal potential of LT-HSCs. Whereas young CD61^Low^ LT-HSCs engrafted perfectly fine in primary recipients, upon secondary transplantation young CD61^Low^ LT-HSCs exhibited a significant decline in their repopulation capacity (Fig. 5F, 5H). Secondary transplantation of aged LT-HSCs revealed that CD61^High^ LT-HSCs retain some self-renewal capacity, whereas aged CD61^Low^ LT-HSCs showed a complete failure in reconstituting the hematopoietic system (Fig. 5G-H).

Even though secondary transplantation of young and aged CD61^High^ and CD61^Low^ LT-HSCs showed differential reconstitution capacity, the contribution to distinct peripheral blood cell populations remained evenly distributed, both in primary and secondary transplantation (Supplementary Fig. S5J - S5U).

As expression of CD150 has been associated with functional activity of HSCs^38^, we assessed whether CD61 expression correlates with expression of CD150. We found no correlation between expression levels of CD61 and CD150 (Supplementary Figure S5V). Thus, differential CD61 expression identifies functionally distinct LT-HSCs populations, independently from CD150 expression.

Collectively, our data show that CD61 marks a functionally superior stem cell population, both in young and aged mice. In aged mice, CD61 marks a population of LT-HSCs that is functionally comparable to young LT-HSCs.

### Repression of CD61 in aged and young LT-HSCs leads to impairment of reconstitution potential

To further assess the role of CD61 we asked whether enforced perturbation of CD61 expression would affect HSC functioning. We repressed CD61 in young and aged LT-HSCs using short-hairpin RNA (shRNAs), with a scrambled shRNA (SCR) as a negative control (Fig. 6A). We isolated LT-HSCs and infected cells ex vivo with a lentiviral pLKO.1 shRNA-mCherry construct. After 5 days of culturing in expansion medium, we transplanted mCherry^+^ cells into lethally irradiated recipients alongside 2×10^6^ WBM W^41^ competitor cells. Downregulation of CD61 was confirmed both on transcriptome and protein level (Fig. 6B-C).

**Figure 6.**
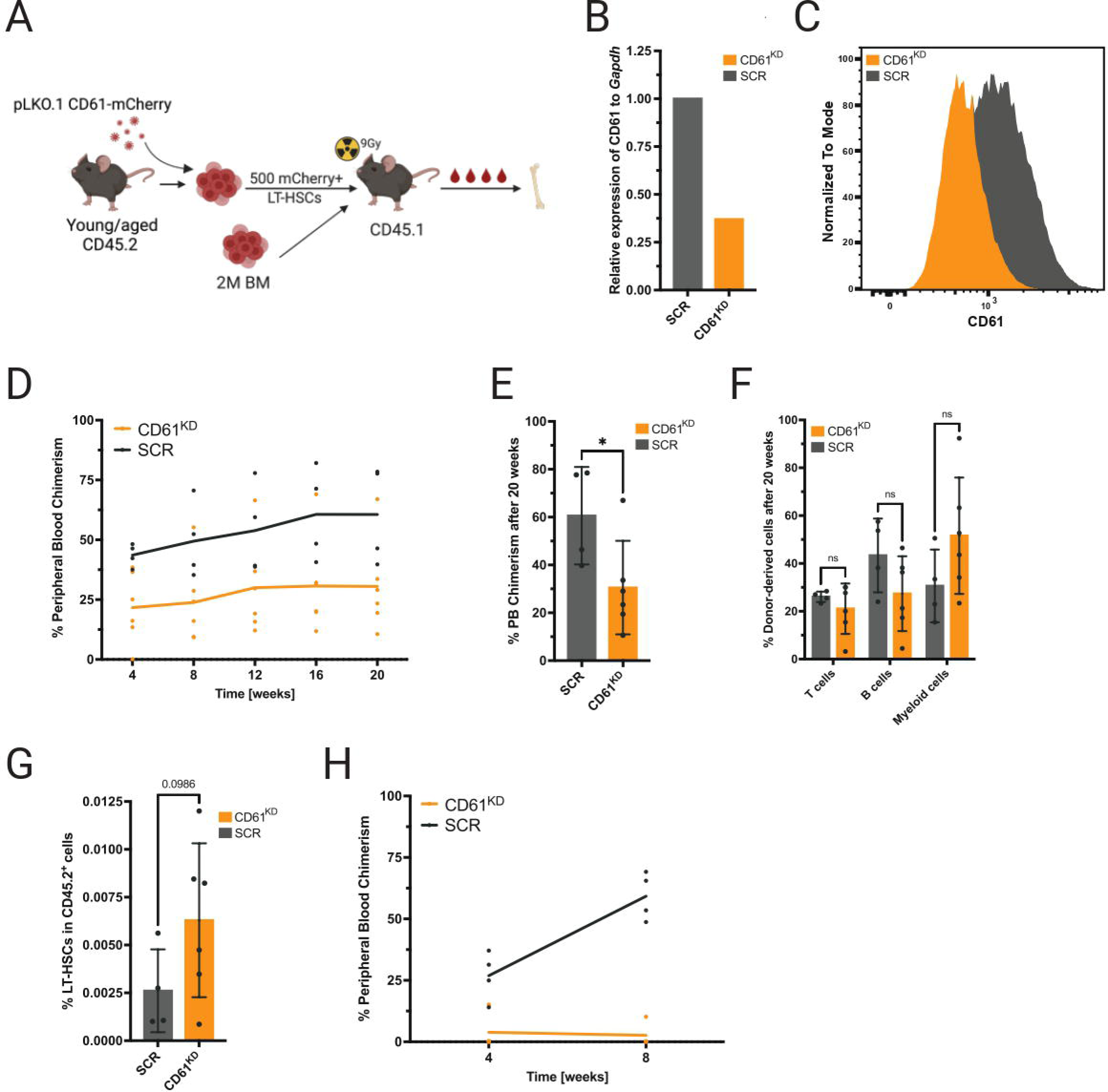
– Transplantation of CD61 repressed LT-HSCs. **A** – Schematic representation of the short-hairpin-mediated CD61 repression **B** – CD61 mRNA levels after shRNA-mediated downregulation of CD61 in aged LT-HSCs **C** – CD61 protein levels after CD61 downregulation, 5 days post-transduction **D** – Competitive transplantation assay of aged LT-HSCs after shRNA mediated CD61 down-regulation (CD61^KD^) compared to scrambled shRNA (SCR) control cells (mice per group – n=4). Data from representative experiment are shown. **E** – Peripheral blood chimerism levels 20 weeks post-transplantation **F** – Donor-derived T-cells, B-cells and myeloid cells, 20 weeks post-transplantation **G** – Donor-derived LT-HSC frequency in bone marrow of recipient mice 20 weeks post-transplantation **H** – Competitive transplantation assay of young LT-HSCs after shRNA mediated CD61 down-regulation (CD61^KD^) compared to scrambled shRNA (SCR) control cells (mice per group – n=4). Data from representative experiment are shown.

Our data revealed that repression of CD61 expression in aged LT-HSCs is detrimental for their reconstitution potential (Fig. 6D, 6E). In contrast to the control that gradually increased their overall contribution to blood production, engraftment of CD61^KD^ LT-HSCs remained stable throughout the whole duration of the experiment. Although the reconstitution potential of CD61^KD^ was limited, the overall ratio of blood cells produced (B, T and myeloid cells) was comparable to that of the SCR control, with a trend toward higher myeloid cells output (Fig. 6F). After 16 weeks the BM of the recipient mice was analyzed. Interestingly, we found a 2-fold increase in the percentage of donor-derived LT-HSCs in CD61^KD^ group, in comparison to the SCR (Fig. 6G). The observed results align closely with the findings from recipient mice transplanted with aged CD61^Low^ LT-HSCs (Fig. 5C, 5E).

Finally, we assessed whether perturbing CD61 expression levels in young LT-HSCs affected functioning as well. Unexpectedly downregulation of CD61 in young LT-HSCs almost completely diminishes their ability to replenish the hematopoietic system in a competitive transplantation experiment (Fig. 6H). Collectively, we show that repression of CD61 expression is detrimental to the engraftment of young and aged LT-HSCs.

## Discussion

In this study we show that differential expression of CD61 identifies transcriptomically and functionally distinct populations of aged LT-HSCs. Notably, we found that LT-HSCs exhibiting high levels of CD61 expression are more quiescent and exhibit superior functional properties compared to LT-HSCs with low CD61 expression. Our findings demonstrate that CD61 is not only a marker that can be used to identify a superior LT-HSC population, but CD61 is also essential for the proper functioning of both young and aged hematopoietic stem cells. Additionally, we provide evidence that aged LT-HSCs expressing high levels of CD61 exhibit a phenotype similar to “young-like” LT-HSCs, suggesting that HSCs displaying increased CD61 expression demonstrate enhanced resilience to the aging process.

The role of CD61 has not been well studied in the context of hematopoietic stem cell aging, although we have recently shown that it is one of the most robustly upregulated genes during aging^23^. It has been shown that young CD61^High^ HSCs display enhanced quiescence and higher repopulation capacity^31^ and our data confirm this. CD61 deficient mice are mostly embryonically lethal, but the few mice that do survive exhibit a phenotype similar to our young CD61^Low^ HSCs. Young CD61 deficient HSCs did not show distinct phenotypic changes in primary transplantation, but lost their self-renewal ability and failed in secondary transplantation, reinforcing the essential role of CD61 expression in HSC self-renewal, possibly by preserving dormancy^29^. It has been reported that while CD61 is dispensable for hematopoiesis, it is required for leukemogenesis^32^.

CD61 expression has been associated with myeloid skewing in aged HSCs^39^, but in our functional experiments we did not observe any myeloid skewing as a result of perturbing CD61 expression. However, we did find a myeloid signature in CD61^High^ LT-HSCs, confirming that, at least on transcriptomic level, CD61^High^ cells might have a myeloid bias^39^.

It remains unknown why CD61 is upregulated in the aged LT-HSC compartment. We observed increased expression of CD61 preferentially in the most primitive compartments, as well as an increase in the population of CD61^High^ LT-HSCs.

Elevated expression of CD61 in primitive LT-HSCs might be caused by age-associated epigenetic alterations, that are known to occur in aged stem cells^15,16,40^. In parallel investigations, we therefore explored whether the epigenetic status of the Itgb3 locus is altered in aged LT-HSCs. However, we have not been able to discern any significant modifications in overall chromatin accessibility, H3K4me3, H3K27me3, H3K36me3, nor DNA methylation patterns between young and aged LT-HSCs (data not shown).

We hypothesize that the age-dependent increase in the population of CD61^High^ LT-HSCs may be the result of a selection process. During aging, dormant CD61^High^ LT-HSCs may preferentially be retained within the aged bone marrow environment, possibly as a result of the many dynamic changes of bone marrow remodeling that occur during aging. Age-associated bone marrow microenvironmental changes involve both cellular as well as non-cellular constituents. One such change is the downregulation of CD61 ligand – Osteopontin, in the aged bone marrow^41^. As a supplementary investigation, we examined the CD61 levels in LT-HSCs from Osteopontin knock-out mice and observed a 2-fold increase in its expression in comparison to WT mice of the same age, suggesting a potential interplay between CD61 and its ligand as a compensatory system in the context of aging (data not shown). This finding highlights the complexity of the interactions between CD61 and its various ligands within the ECM during aging and might explain the favored CD61^High^ expression in aged HSCs.

Our data also show that CD61^High^ LT-HSCs are quiescent and functionally superior, and that inhibition of CD61-CD51 heterodimerization induces cell cycling of CD61^High^ HSCs. Upon aging, alterations in the bone marrow niche may cause stress to the HSCs residing in it, and we propose that CD61-CD51-mediated maintenance of HSCs dormancy is one way for the stem cells to retain their repopulating potential and protect themselves against the detrimental changes of the environment.

Finally, our study reveals that highly purified LT-HSCs can be further fractionated based on differential CD61 expression into populations of cells with distinct repopulating ability. Notably, a small population of aged CD61^High^ HSCs demonstrated comparable stem cell potential and self-renewal capacity to their young counterparts. This finding supports the existence and allows the prospective isolation of “young-like” stem cells within an aged stem cell compartment. As it has been well demonstrated that dormant HSCs engraft and repopulate better than cycling HSCs^34^, we assume that CD61-mediated quiescence directly contributes to repopulating potential. In addition, it is likely that CD61^High^ LT-HSCs have accumulated less replicative stress (supported also by lower levels of DNA damage), and are functionally younger and more potent. If our findings can be corroborated for human HSCs, this may open avenues to prospectively isolate functionally superior stem cells for clinical transplantations.

## Acknowledgment

We thank UMCG Flowcytometry Unit facility staff J. Teunis, T. Bijma and G. Mesander for their assistance on cell sorting and analysis. We also are very grateful to dr. L. Bystrykh at ERIBA for assistance with data analysis and A. Svendsen for sharing the epigenetic data.

This work was supported by the ARCH (a European Union’s Horizon 2020 Research and Innovation Program) under Marie Skłodowska-Curie grant agreement 813091 and by the Netherlands Organization for Scientific Research (NWO) grant nr. 12583.

## Authorship contribution

Contribution: N.S. and G.d.H. conceptualized the study; N. S. and G.d.H carried out the methodology; N.S, I.S.F., A.D-A. and E.W. performed the investigation; N.S. performed the formal analysis; N.S. curated the data; N.S. wrote the original manuscript draft; N.S and G.d.H. reviewed and edited the manuscript; G.d.H. supervised the study.

Conflict-of-interest disclosure: The authors declare no competing financial interests.

## References

1. De Haan G, Lazare SS. Aging of hematopoietic stem cells. Blood. 2018;131(5):479–487. doi:10.1182/blood-2017-06-746412

2. Mejia-Ramirez E, Florian MC. Understanding intrinsic hematopoietic stem cell aging. Haematologica. 2020;105(1):22–37. doi:10.3324/haematol.2018.211342

3. Ho YH, Méndez-Ferrer S. Microenvironmental contributions to hematopoietic stem cell aging. Haematologica. 2020;105(1):38–46. doi:10.3324/haematol.2018.211334

4. Dykstra B, Olthof S, Schreuder J, Ritsema M, Haan G De. Clonal analysis reveals multiple functional defects of aged murine hematopoietic stem cells. J Exp Med. 2011;208(13):2691–2703. doi:10.1084/jem.20111490

5. Beerman I, Bhattacharya D, Zandi S, et al. Functionally distinct hematopoietic stem cells modulate hematopoietic lineage potential during aging by a mechanism of clonal expansion. Proc Natl Acad Sci U S A. 2010;107(12):5465–5470. doi:10.1073/pnas.1000834107

6. Yamamoto R, Wilkinson AC, Ooehara J, et al. Large-Scale Clonal Analysis Resolves Aging of the Mouse Hematopoietic Stem Cell Compartment. Cell Stem Cell. 2018;22(4):600–607.e4. doi:10.1016/j.stem.2018.03.013

7. Chambers SM, Shaw CA, Gatza C, Fisk CJ, Donehower LA, Goodell MA. Aging hematopoietic stem cells decline in function and exhibit epigenetic dysregulation. PLoS Biol. 2007;5(8):1750–1762. doi:10.1371/journal.pbio.0050201

8. Rossi DJ, Bryder D, Zahn JM, et al. Cell intrinsic alterations underlie hematopoietic stem cell aging. Proc Natl Acad Sci U S A. 2005;102(26):9194–9199. doi:10.1073/pnas.0503280102

9. Florian MC, Dörr K, Niebel A, et al. Cdc42 activity regulates hematopoietic stem cell aging and rejuvenation. Cell Stem Cell. 2012;10(5):520–530. doi:10.1016/j.stem.2012.04.007

10. Amoah A, Keller A, Emini R, et al. Aging of human hematopoietic stem cells is linked to changes in Cdc42 activity. Haematologica. 2022;107(2):393–402. doi:10.3324/haematol.2020.269670

11. Ho TT, Warr MR, Adelman ER, et al. Autophagy maintains the metabolism and function of young and old stem cells. Nature. 2017;543(7644):205-210. doi:10.1038/nature21388

12. Dong S, Wang Q, Kao YR, et al. Chaperone-mediated autophagy sustains haematopoietic stem-cell function. Nature. 2021;591(7848):117-123. doi:10.1038/s41586-020-03129-z

13. Vas V, Senger K, Dörr K, Niebel A, Geiger H. Aging of the microenvironment influences clonality in hematopoiesis. PLoS One. 2012;7(8):1–6. doi:10.1371/journal.pone.0042080

14. Walter D, Lier A, Geiselhart A, et al. Exit from dormancy provokes DNA-damage-induced attrition in haematopoietic stem cells. Nature. 2015;520(7548):549-552. doi:10.1038/nature14131

15. Beerman I, Bock C, Garrison BS, et al. Proliferation-dependent alterations of the DNA methylation landscape underlie hematopoietic stem cell aging. Cell Stem Cell. 2013;12(4):413–425. doi:10.1016/j.stem.2013.01.017

16. Sun D, Luo M, Jeong M, et al. Epigenomic profiling of young and aged HSCs reveals concerted changes during aging that reinforce self-renewal. Cell Stem Cell. 2014;14(5):673–688. doi:10.1016/j.stem.2014.03.002

17. Ergen A V., Boles NC, Goodell MA. Rantes/Ccl5 influences hematopoietic stem cell subtypes and causes myeloid skewing. Blood. 2012;119(11):2500–2509. doi:10.1182/blood-2011-11-391730

18. Ho YH, del Toro R, Rivera-Torres J, et al. Remodeling of Bone Marrow Hematopoietic Stem Cell Niches Promotes Myeloid Cell Expansion during Premature or Physiological Aging. Cell Stem Cell. 2019;25(3):407–418.e6. doi:10.1016/j.stem.2019.06.007

19. Saçma M, Pospiech J, Bogeska R, et al. Haematopoietic stem cells in perisinusoidal niches are protected from ageing. Nat Cell Biol. 2019;21(11):1309–1320. doi:10.1038/s41556-019-0418-y

20. Verovskaya E, Broekhuis MJC, Zwart E, et al. Heterogeneity of young and aged murine hematopoietic stem cells revealed by quantitative clonal analysis using cellular barcoding. Blood. 2013;122(4):523–532. doi:10.1182/blood-2013-01- 481135

21. Grover A, Sanjuan-Pla A, Thongjuea S, et al. Single-cell RNA sequencing reveals molecular and functional platelet bias of aged haematopoietic stem cells. Nat Commun. 2016;7. doi:10.1038/ncomms11075

22. Wilson NK, Kent DG, Buettner F, et al. Combined Single-Cell Functional and Gene Expression Analysis Resolves Heterogeneity within Stem Cell Populations. Cell Stem Cell. 2015;16(6):712–724. doi:10.1016/j.stem.2015.04.004

23. Flohr Svendsen A, Yang D, Kim KM, et al. A comprehensive transcriptome signature of murine hematopoietic stem cell aging. Blood. 2021;138(6):439–451. doi:10.1182/blood.2020009729

24. Prosper F, Verfaillie CM. Regulation of hematopoiesis through adhesion receptors. J Leukoc Biol. 2001;69(3):307–316. doi:10.1189/jlb.69.3.307

25. Shattil SJ, Kim C, Ginsberg MH. The final steps of integrin activation: The end game. Nat Rev Mol Cell Biol. 2010;11(4):288–300. doi:10.1038/nrm2871

26. Antonov AS, Kolodgie FD, Munn DH, Gerrity RG. Regulation of macrophage foam cell formation by αVβ3 integrin: Potential role in human atherosclerosis. Am J Pathol. 2004;165(1):247–258. doi:10.1016/S0002-9440(10)63293-2

27. Rapisarda V, Borghesan M, Miguela V, et al. Integrin Beta 3 Regulates Cellular Senescence by Activating the TGF-β Pathway. Cell Rep. 2017;18(10):2480–2493. doi:10.1016/j.celrep.2017.02.012

28. Bachmann M, Kukkurainen S, Hytönen VP, Wehrle-Haller B. Cell adhesion by integrins. Physiol Rev. 2019;99(4):1655–1699. doi:10.1152/physrev.00036.2018

29. Umemoto T, Yamato M, Ishihara J, et al. Integrin-αvβ3 regulates thrombopoietin-mediated maintenance of hematopoietic stem cells. Blood. 2012;119(1):83–94. doi:10.1182/blood-2011-02-335430

30. Umemoto T, Matsuzaki Y, Shiratsuchi Y, et al. Integrin αvβ3 enhances the suppressive effect of interferon-γ on hematopoietic stem cells. EMBO J. 2017;36(16):2390–2403. doi:10.15252/embj.201796771

31. Umemoto T, Yamato M, Shiratsuchi Y, et al. Expression of Integrin β 3 Is Correlated to the Properties of Quiescent Hemopoietic Stem Cells Possessing the Side Population Phenotype . J Immunol. 2006;177(11):7733–7739. doi:10.4049/jimmunol.177.11.7733

32. Miller PG, Al-Shahrour F, Hartwell KA, et al. InVivo RNAi Screening Identifies a Leukemia-Specific Dependence on Integrin Beta 3 Signaling. Cancer Cell. 2013;24(1):45–58. doi:10.1016/j.ccr.2013.05.004

33. Rodriguez-Fraticelli AE, Weinreb C, Wang SW, et al. Single-cell lineage tracing unveils a role for TCF15 in haematopoiesis. Nature. 2020;583(7817):585-589. doi:10.1038/s41586-020-2503-6

34. Cabezas-Wallscheid N, Buettner F, Sommerkamp P, et al. Vitamin A-Retinoic Acid Signaling Regulates Hematopoietic Stem Cell Dormancy. Cell. 2017;169(5):807–823.e19. doi:10.1016/j.cell.2017.04.018

35. Hynes R. Integrins: Bidirectional, allosteric signaling machines. Cell. 2002;110:673–687. doi:10.1016/s0092-8674(02)00971-6

36. Hartman GD, Egbertson MS, Halczenko W, et al. Non-Peptide Fibrinogen Receptor Antagonists. 1. Discovery and Design of Exosite Inhibitors’. J. Med. Chem. 1992, 35, 24, 4640–4642. doi: 10.1021/jm00102a020

37. Haubner R, Wester H-J, Burkhart F, et al. Glycosylated RGD-Containing Peptides: Tracer for Tumor Targeting and Angiogenesis Imaging with Improved Biokinetics. J Nucl Med. 2001 Feb;42(2):326–36.

38. Kiel MJ, Yilmaz ÖH, Iwashita T, Yilmaz OH, Terhorst C, Morrison SJ. SLAM family receptors distinguish hematopoietic stem and progenitor cells and reveal endothelial niches for stem cells. Cell. 2005;121(7):1109–1121. doi:10.1016/j.cell.2005.05.026

39. Mann M, Mehta A, de Boer CG, et al. Heterogeneous Responses of Hematopoietic Stem Cells to Inflammatory Stimuli Are Altered with Age. Cell Rep. 2018;25(11):2992–3005.e5. doi:10.1016/j.celrep.2018.11.056

40. Wahlestedt M, Norddahl GL, Sten G, et al. An epigenetic component of hematopoietic stem cell aging amenable to reprogramming into a young state. Blood. 2013;121(21):4257–4264. doi:10.1182/blood-2012-11-469080

41. Guidi N, Sacma M, Ständker L, et al. Osteopontin attenuates aging-associated phenotypes of hematopoietic stem cells. EMBO J. 2017;36(7):840–853. doi:10.15252/embj.201694969

