## Supplementary_figures1-6 for "CD61 identifies a superior population of aged murine hematopoietic stem cells and is required to preserve quiescence and self-renewal"

### Supplementary Figure 1

## A

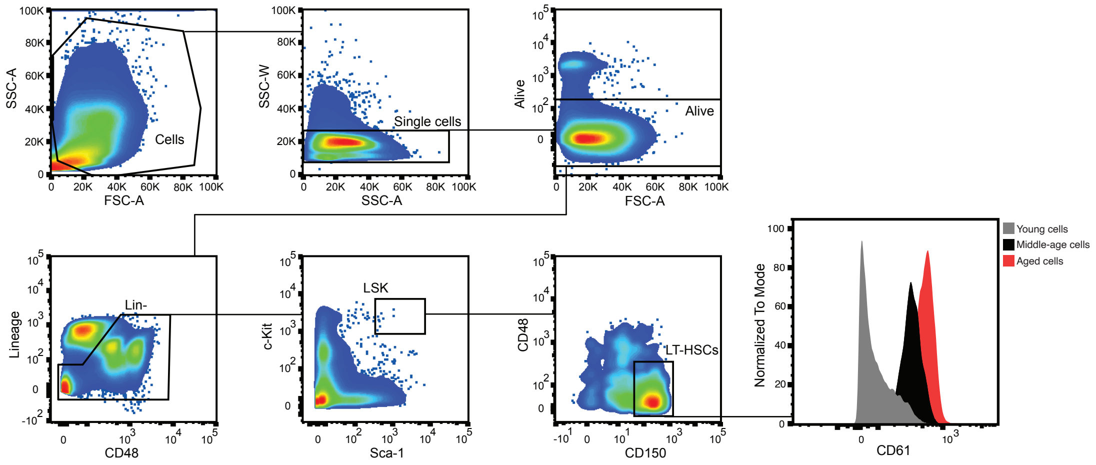

### Supplementary Figure 2

A

Myeloid signature
(Mann, M. et al, 2018)

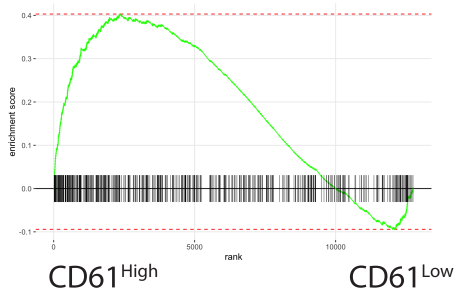


B

Aged HSC signature
(Svendsen, A. et al, 2021)

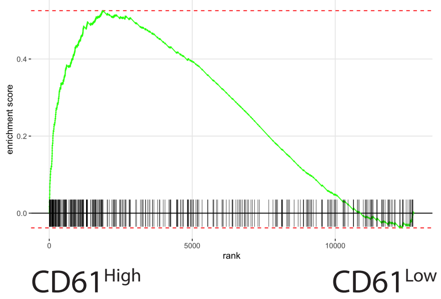


C

List of Aging Signature genes
found in the CD61 DEG

| |
|---|
| Alcam |
| Fap |
| Bmpr1a |
| Cd34 |
| Egr1 |
| Il12rb2 |
| C4b |
| Gstm1 |
| Gda |
| Mlec |
| Maf |
| Tbc1d8 |
| Ldhd |
| Cd200r4 |

D

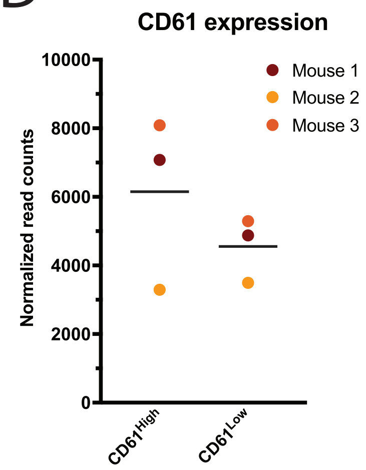

### Supplementary Figure 3

A

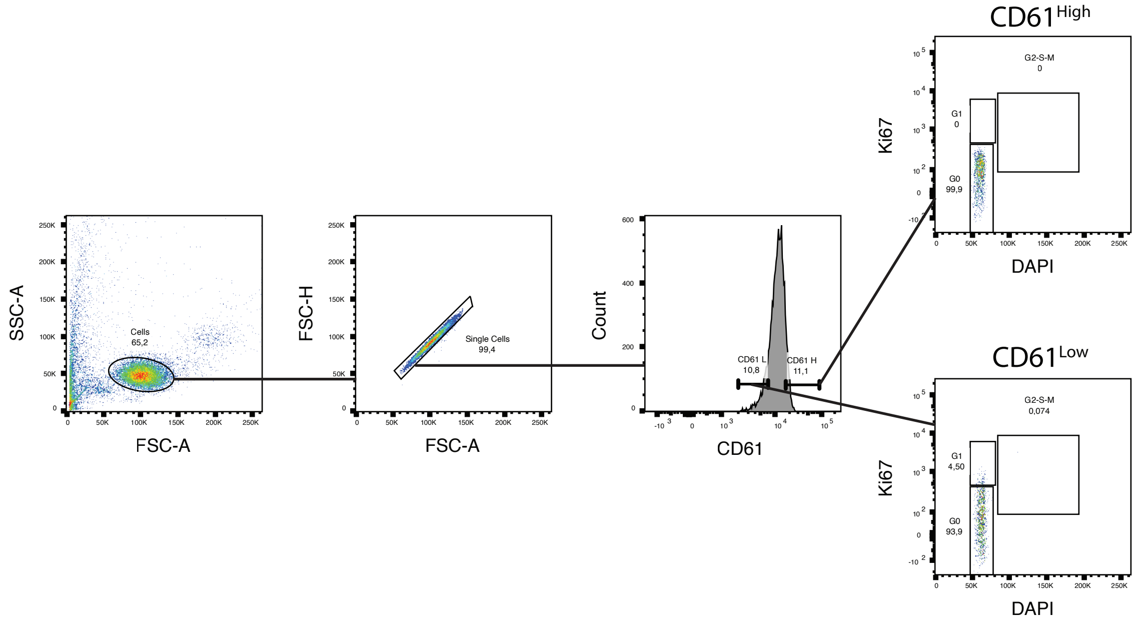

### Supplementary Figure 4

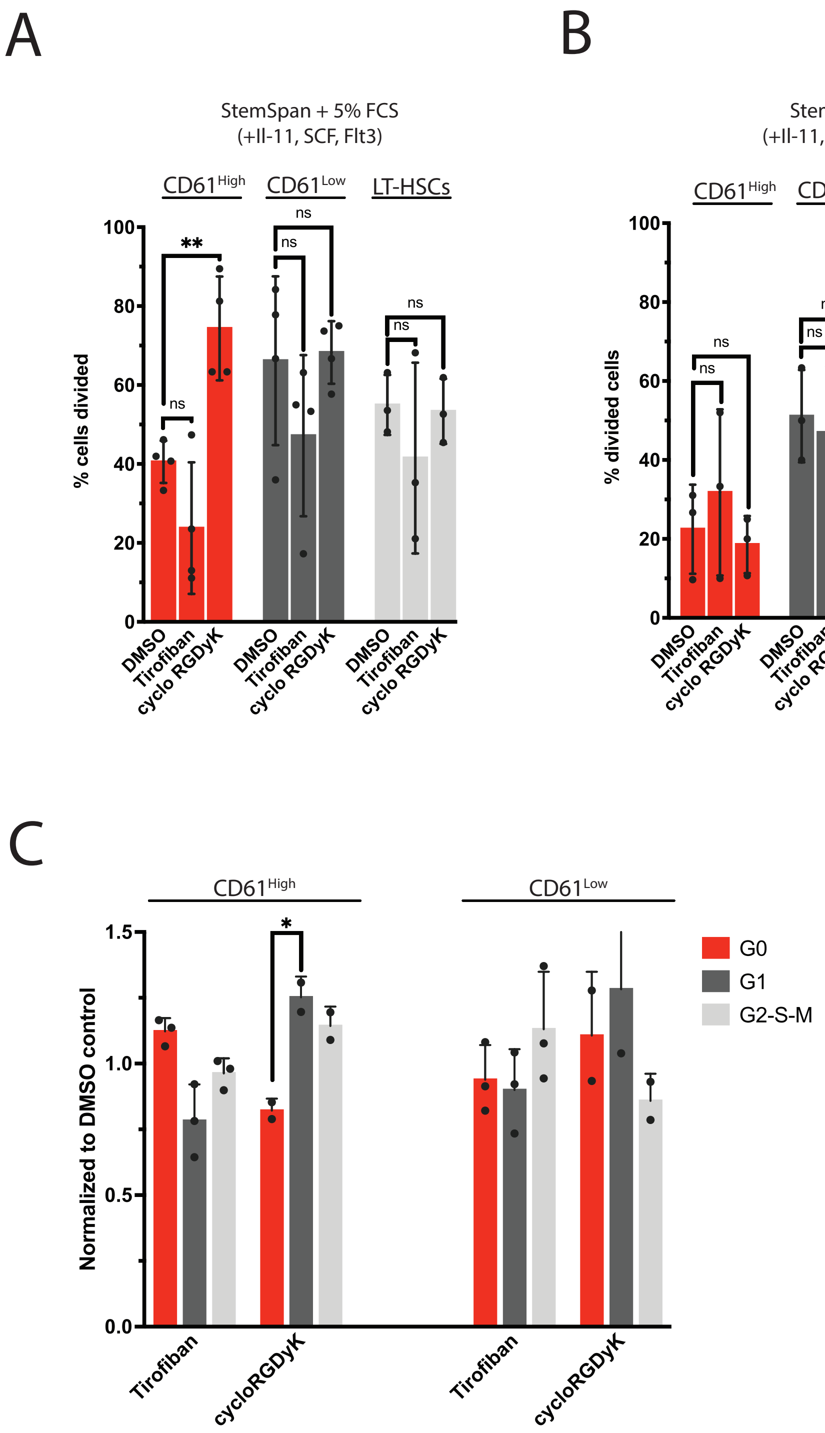

### Supplementary Figure 5

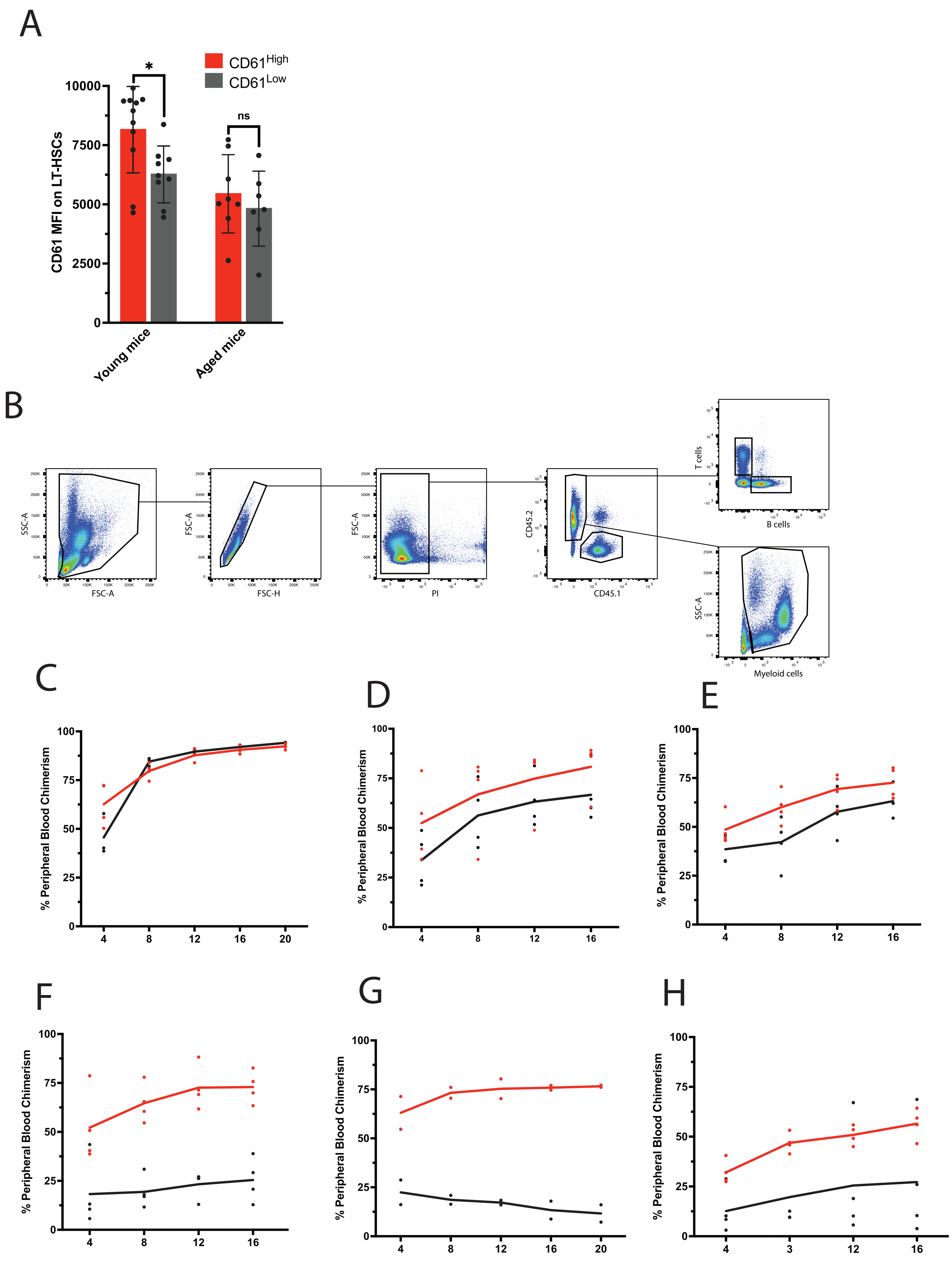

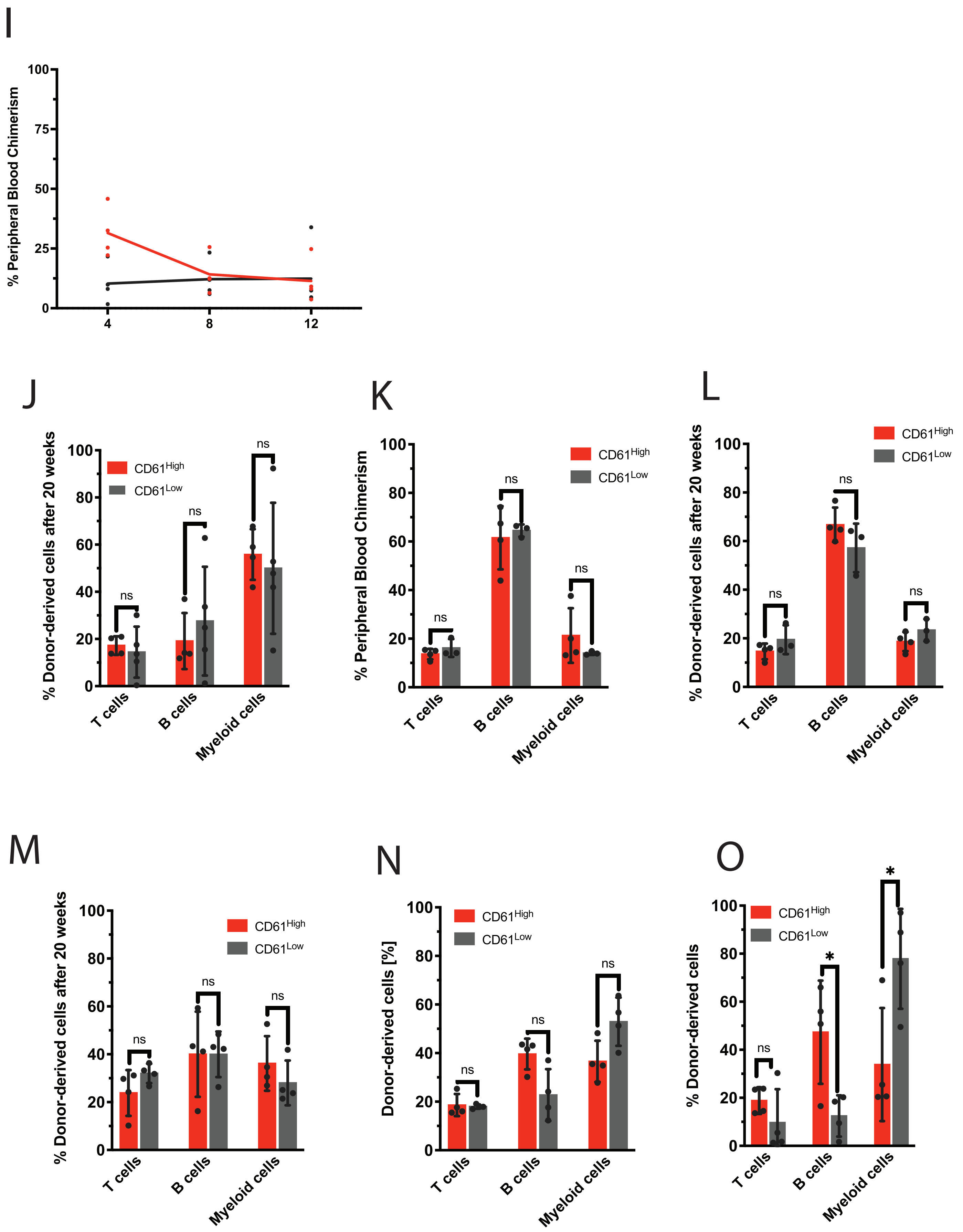
I
% Peripheral Blood Chimerism
100
75
50
25
0
4
8
12
J
% Donor-derived cells after 20 weeks
CD61High
CD61Low
ns
T cells
B cells
Myeloid cells
K
% Peripheral Blood Chimerism
CD61High
CD61Low
ns
T cells
B cells
Myeloid cells
L
% Donor-derived cells after 20 weeks
CD61High
CD61Low
ns
T cells
B cells
Myeloid cells
M
% Donor-derived cells after 20 weeks
CD61High
CD61Low
ns
T cells
B cells
Myeloid cells
N
Donor-derived cells [%]
CD61High
CD61Low
ns
T cells
B cells
Myeloid cells
O
% Donor-derived cells
CD61High
CD61Low
ns
*
T cells
B cells
Myeloid cells

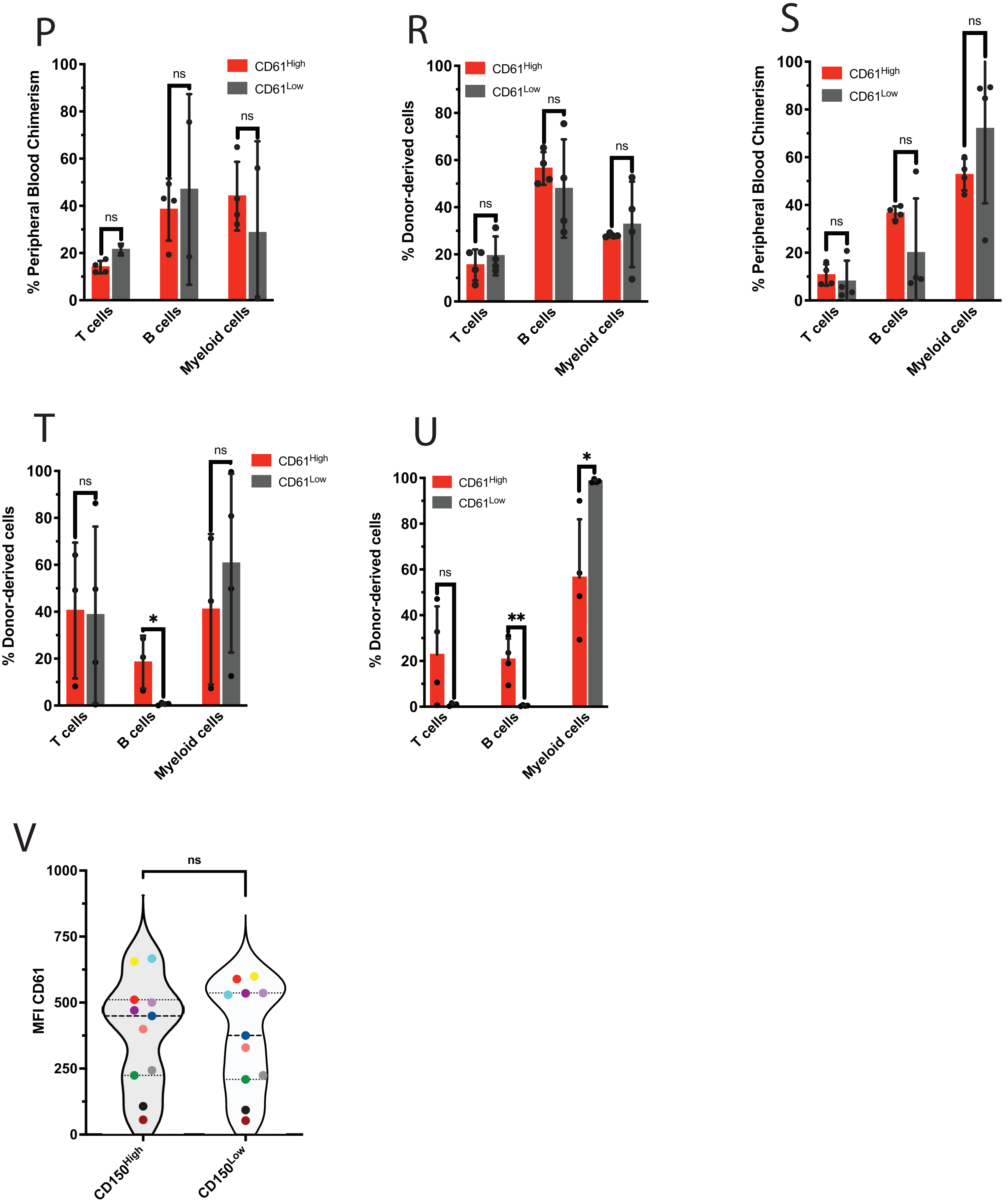

P
% Peripheral Blood Chimerism
CD61High
CD61Low
ns
T cells
B cells
Myeloid cells
R
% Donor-derived cells
S
T
*
U
**
V
MFI CD61
CD150High
CD150Low
0
20
40
60
80
100
250
500
750
1000

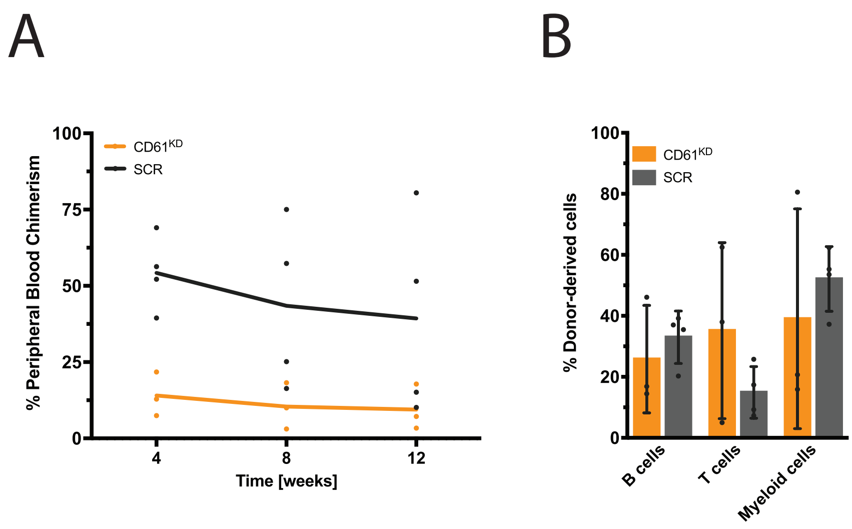
A
B
% Peripheral Blood Chimerism
100
75
50
25
0
4
8
12
Time [weeks]
CD61KD
SCR
% Donor-derived cells
100
80
60
40
20
0
B cells
T cells
Myeloid cells
CD61KD
SCR
