## Supplementary_fig_legends for "CD61 identifies a superior population of aged murine hematopoietic stem cells and is required to preserve quiescence and self-renewal"

**Supplementary Figure Legends**

**Supplementary Figure 1**

**A -** Sorting strategy for CD61^High^ and CD61^Low^ LT-HSCs from young, middle-aged and aged mice

**Supplementary Figure 2**

**A -** Signature enrichment plot from GSEA using Myeloid Signature gene set in LT-HSCs from CD61^High^ and CD61^Low^ groups (GSE100428)

**B -** Signature enrichment plot from GSEA using Aging Signature gene set in LT-HSCs from CD61^High^ and CD61^Low^ groups

**C –** List of Aging Signature genes present in the CD61 DEG

**D –** Normalized read counts for CD61 in CD61^High^ and CD61^Low^ groups

**Supplementary Figure 3**

**A –** LT-HSC analysis strategy for cell cycle analysis

**Supplementary Figure 4**

**A -** Single cell division assay of aged CD61^High^, CD61^Low^ and unfractionated LT-HSCs treated with DMSO only, Tirofiban or cycloRGDγK in StemSpan with 5% serum supplementation and cytokines

**B -** Single cell division assay of aged CD61^High^, CD61^Low^ and unfractionated LT-HSCs treated with DMSO only, Tirofiban of cycloRGDγK in serum-free StemSpan supplemented with cytokines

**C -** Cell cycle analysis using Ki67/DAPI staining of young CD61^High^ and CD61^Low^ LT-HSCs treated with Tirofiban or cycloRGDγK in StemSpan; The results are normalized to DMSO control

**Supplementary Figure 5**

**A –** CD61 MFI in donor-derived LT-HSCs from primary recipient BM of young and aged donor LT-HSCs

**B –** Representative analysis of PB by flow cytometry

**C -** Competitive transplantation assay of 500 CD61^High^ and CD61^Low^ LT-HSCs from young mice into lethally irradiated recipients (mice per group – n=4); 2^nd^ replicate

**D -** Competitive transplantation assay of 500 CD61^High^ and CD61^Low^ LT-HSCs from young mice into lethally irradiated recipients (mice per group – n=4); 3^rd^ replicate

**E -** Secondary transplantation assay of 500 LT-HSCs derived from primary recipients transplanted with young CD61^High^ and CD61^Low^ LT-HSCs (mice per group – n=4); 2^nd^ replicate

**F -** Secondary transplantation assay of 500 LT-HSCs derived from primary recipients transplanted with young CD61^High^ and CD61^Low^ LT-HSCs (mice per group – n=4); 3^rd^ replicate

**G –** Competitive transplantation assay of 500 CD61^High^ and CD61^Low^ LT-HSCs from aged mice into lethally irradiated recipients (mice per group – n=4); 2^nd^ replicate

**H –** Competitive transplantation assay of 500 CD61^High^ and CD61^Low^ LT-HSCs from aged mice into lethally irradiated recipients (mice per group – n=4); 3^rd^ replicate

**I –** Secondary transplantation assay of 500 LT-HSCs derived from primary recipients transplanted with aged CD61^High^ and CD61^Low^ LT-HSCs (mice per group – n=4); 2^nd^ replicate

**J –** Donor-derived PB cell populations post-transplantation from primary young CD61^High^ and CD61^Low^ LT-HSCs transplantation;1^st^ replicate

**K –** Donor-derived PB cell populations post-transplantation from primary young CD61^High^ and CD61^Low^ LT-HSCs transplantation;2^nd^ replicate

**L –** Donor-derived PB cell populations post-transplantation from primary young CD61^High^ and CD61^Low^ LT-HSCs transplantation;3^rd^ replicate

**M –** Donor-derived PB cell populations post-transplantation from secondary young CD61^High^ and CD61^Low^ LT-HSCs transplantation;1^st^ replicate

**N –** Donor-derived PB cell populations post-transplantation from secondary young CD61^High^ and CD61^Low^ LT-HSCs transplantation; 2^nd^ replicate

**O –** Donor-derived PB cell populations post-transplantation from secondary young CD61^High^ and CD61^Low^ LT-HSCs transplantation;3^rd^ replicate

**P –** Donor-derived PB cell populations post-transplantation from primary aged CD61^High^ and CD61^Low^ LT-HSCs transplantation;1^st^ replicate

**R –** Donor-derived PB cell populations post-transplantation from primary aged CD61^High^ and CD61^Low^ LT-HSCs transplantation;2^nd^ replicate

**S –** Donor-derived PB cell populations post-transplantation from primary aged CD61^High^ and CD61^Low^ LT-HSCs transplantation;3^rd^ replicate

**T –** Donor-derived PB cell populations post-transplantation from secondary aged CD61^High^ and CD61^Low^ LT-HSCs transplantation;1^st^ replicate

**U –** Donor-derived PB cell populations post-transplantation from secondary aged CD61^High^ and CD61^Low^ LT-HSCs transplantation; 2^nd^ replicate

**V –** CD61 MFI levels in the 10% highest and 10% lowest CD150-expressing cells

**Supplementary Figure 6**

**A –** Competitive transplantation assay of 500 CD61^KD^ and SCR LT-HSCs isolated from aged mice into lethally irradiated recipients (mice per group – n=4); 2^nd^ replicate

**B -** Donor-derived PB cell populations post-transplantation from aged CD61^KD^ and SCR LT-HSCs transplantation; 2^nd^ replicate
