## Supplementary_material for "CD61 identifies a superior population of aged murine hematopoietic stem cells and is required to preserve quiescence and self-renewal"

**Supplementary Materials and Methods**

**Mice**

All experiments were approved by the Central Commission for animal Testing and Animal Ethical Committee. C57BL/6.SJL (10-16 weeks) and C57BL/6J-kitW41/kitW-41J (W41) were obtained from the Central Animal Facility (CDP) at the University Medical Center Groningen. Young (2-5 months) and aged (>22 months) C57BL/6J were obtained from either CDP or Janvier labs, France. All mice were housed in a temperature and day cycle-controlled conditions.

**Flow Cytometry**

*HSC and progenitor cells isolation*

Bone marrow was isolated from the tibia, femur, pelvis, sternum and spine by crushing and the erythrocytes were lysed with erylysis buffer (17mM NaCl, 0.16M NH_4_Cl, 0.13mM EDTA). For LT-HSC and progenitor cells isolation, the lysed bone marrow was stained with c-Kit-PE/BB515, Sca-1-BV421, CD48-Alexa Fluor 647, CD150-PE-Cy7 and lineage (Lin) marker antibodies (B220, CD3, Gr-1, Mac-1 and Ter-119) conjugated with Alexa Fluor 700. After incubation with the antibodies for 30min at 4°C, the sample was washed and filtered. PI was used for Live/Dead staining. The LT-HSCs (Lin^-^, c-Kit^+^, Sca-1^+^, CD48^-^, CD150^+^), ST-HSCs (Lin^-^, c-Kit^+^, Sca-1^+^, CD48^-^, CD150^-^) and MPPs (Lin^-^, c-Kit^+^, Sca-1^+^, CD48^+^, CD150^-^) were isolated on MoFlo Astrios or MoFlo XDP cell sorters (Beckman Coulter). For CD61, CD61-PE antibody was used.

**In vitro experiments**

*Single cell colony assay*

Single LT-HSCs, isolated from young or aged mouse, were sorted into a 96-well round-bottom plate and cultured for 14 days in StemSpan (STEMCELL Technologies) supplemented with 100U/ml penicillin, 100μg/ml streptomycin (Gibco), 10% Australian FCS, 300ng/ml SCF (Peprotech), 20ng/ml IL-11 (R&D Systems) and 1ng/ml Flt3 ligand (Peprotech). The size of the colonies was analyzed after 7 and 14 days. Size 0: no cells; Size 1: 1-30; Size 2: 31-100 cells; Size 3: 101-1.000; Size 4: 5.000 cells; Size 5: 15.000 cells; Size 6: 30.000 cells; Size 7: >30.000

*Single cell division assay*

Single LT-HSCs, isolated from young or aged mouse, were sorted into a 60-well Terasaki plate and cultured for 2 days in StemSpan (STEMCELL Technologies) supplemented with 100U/ml penicillin, 100μg/ml streptomycin (Gibco), 10% Australian FCS, 300ng/ml SCF (Peprotech), 20ng/ml IL-11 (R&D Systems) and 1ng/ml Flt3 ligand (Peprotech). The number of cells was analyzed after 48h and categorized as 1 - 1 division; >2 – more than 2 cells.

*γH2AX immunofluorescence staining*

1.000-2.000 LT-HSCs were sorted directly onto spots of an adhesion slide (VWR) filled with PBS. Cells were then permeabilized with 4% PFA and permeabilized with 0.25% Triton X-100 in PBS and blocked with 5% BSA + 0.05% Tween-20 in PBS, at RT and subsequently stained with 1:400 α-γH2AX rabbit antibody ON at 4°C. Cells were stained with 1:500 secondary antibody goat-anti-rabbit Alexa Fluor 488 and the coverslip was mounted with ProLong Antifade Mounant with DAPI (Thermo Fisher). The slide was imaged on a Leica Sp8 confocal microscope. The data was analyzed using Fiji Image J.

*LT-HSC cell cycle analysis*

10.000-20.000 LT-HSCs were sorted and fixed and stained according to the manufacturer’s protocol (Fix/Perm kit #554714). Cells were stained with 1:200 Ki67-PE-Cy7 antibody and 2μg/ml DAPI. Samples were analyzed on BD FACS Canto II or BD Symphony.

*Inhibitor treatment of LT-HSCs*

For single-cell proliferation assay and division assay, aged LT-HSCs were sorted directly into wells with medium (described above) and supplemented with 50ng/mL cycloRGDγK (Selleckchem, S7844) and 25ng/mL Tirofiban (Selleckchem, S3085).

For cell cycle analysis, at least 10.000 freshly isolated LT-HSCs were plated in 12-well plate in medium (described above), supplemented with 50ng/mL cycloRGDγK and 25ng/mL Tirofiban and incubated for 2h in 37°C, then the protocol for cell cycle analysis was followed.

For single-cell expansion assay aged LT-HSCs were sorted directly into rond-bottom 96-wells plate with HemEx-Type9A (Nipro) medium supplemented with 100ng/ml TPO, 10ng/ml SCF, 50ng/mL cycloRGDγK (Selleckchem, S7844) and 25ng/mL Tirofiban (Selleckchem, S3085). After 14 days the size of the colonies was analyzed based on the scoring range of single-cell proliferation assay.

*Quantitative PCR analysis*

5.000 or more LT-HSCs, ST-HSCs or MPPs were sorted directly into RLT Plus lysis buffer and the RNA extraction was carried out using RNeasy Micro Kit (Qiagen). The RNA was then transcribed into cDNA using SuperScript VILO cDNA Synthesis Kit (Invitrogen). The amplicons for CD61 and housekeeping gene GAPDH were amplified and quantified via qPCR using LightCycler SYBR Green I MasterMix and LightCycler 480 Instrument (Roche). CD61 expression was normalized relative to the housekeeping gene.

**Transplantation**

*CD61^High^ and CD61^Low^ LT-HSC transplantation*

500 CD61^High^ and CD61^Low^ LT-HSCs were sorted from young or aged CD45.2^+^ C57BL/6 mouse and were transplanted together with 2 million CD45.1^+^ W41 whole bone marrow cells into gender-matched, lethally irradiated (9 Gy) CD45.1^+^ C57BL/6.SJL recipients, respectively. 16 weeks after transplantation, 500 CD45.2^+^ donor-derived LT-HSCs were isolated from the primary recipients of CD61^High^ and CD61^Low^ groups and transplanted alongside 2 million CD45.1^+^ W41 bone marrow cells into gender-matched lethally irradiated (9 Gy) CD45.1^+^ C57BL/6.SJL secondary recipients.

*CD61^KD^ LT-HSC transplantation*

**CD61^KD^** Oligonucleotides for CD61^KD^ were annealed and cloned into the empty pLKO.1_mCherry vector upon digestion with AgeI and EcoRI. The virus was produced by transfecting LentiX HEK293 cells with the construct alongside packaging vectors (pCMV8.91, VSV-G) using FuGENE.

**Transduction** After 24h the medium was changed to HemEx-Type9A and the viral supernatant was collected 24h later. LT-HSCs were isolated from CD45.2^+^ C57BL/6 mouse and plated on retronectin-coated plates 24h prior to transduction in HemEx-Type9A (Nipro) medium supplemented with 100ng/ml TPO and 10ng/ml SCF. The medium was removed and the filtered viral supernatant added to the LT-HSCs. The plate was centrifuged for 45min at 400g and incubated for 5 days at 37°C.

**Transplantation** After 5 days the cells were collected from the wells and 500 GFP^+^/mCherry^+^ LT-HSCs were sorted and transplanted alongside 2 million CD45.1^+^ W41 bone marrow cells into gender-matched lethally irradiated (9 Gy) CD45.1^+^ C57BL/6.SJL recipients.

| Name | Sequence |
| --- | --- |
| shRNA.1 | CCGG GCATCCCATTTGCTAGTGTTT CTCGAG AAACACTAGCAAATGGGATGCTTTTT |
| shRNA.2 | CCGG GCTCATCTGGAAGCTACTCAT CTCGAG ATGAGTAGCTTCCAGATGAGCTTTTT |
| shRNA.3 | CCGG CGCCGTGAATTGTACCTACAA CTCGAG TTGTAGGTACAATTCACGGCGTTTTT |

*Peripheral blood count*

Peripheral blood was collected from the retro-orbital venous plexus in heparinized capillary tubes. 25 μl PB was used for cell counting using Medonic CA-620.

**RNA-sequencing**

Total RNA was isolated from 5000 LT-HSCs using RNeasy Plus Micro Kit (QIAGEN, 74034) according to the manufacturer’s instructions. Library preparation was performed with SMART Ultra-Low Input kit v4 (Takara) and Nextera™ XT Library Prep kit (Illumina). Samples were sequenced using a NextSeq 5000 (Illumina) in the same flow cell and were pulled equimolarly.

**Data analysis** FASTQ files were quality-control checked using FastQC (0.11.9) and Picard (2.23.0). Following, reads were mapped to reference mouse genome (GENCODE, GRCm38, M13) using STAR (2.7.0d) with standard arguments and --outFilterMatchNminOverLread 0.4 --outFilterScoreMinOverLread 0.4. Unstranded read counts were used to perform differential expression analysis using DESeq2.

**Statistical analysis**

All experiments were performed as least 2 times. The number of mice or technical replicates is indicated in the figure legends. Data are shown as mean ± SD. Unpaired, two-tailed Student t-test and two-way ANOVA with Sidak’s multiple comparison test were performed in GraphPad Prism 9.0 and 10.0. Significant p-value was indicated as *p<0.05, **p<0.01, ***p<0.001 and ****p<0.0001
